# Global nonequilibrium cortical dynamics tie mid-level pupil-linked arousal to optimal task performance in humans

**DOI:** 10.64898/2026.06.02.729595

**Authors:** Elvira del Agua, Stijn Adriaan Nuiten, Jasper B. Zantvoord, Gustavo Deco, Simon van Gaal

**Affiliations:** Faculty of Medicine and Life Sciences, Center for Brain and Cognition, Universitat Pompeu Fabra, Barcelona, Spain; University Psychiatric Clinics (UPK), University of Basel, Basel, Switzerland; Amsterdam UMC location University of Amsterdam, Department of Psychiatry, Amsterdam, The Netherlands; Amsterdam Neuroscience, Amsterdam, The Netherlands; Institució Catalana de la Recerca i Estudis Avançats (ICREA), Barcelona, Spain; International Centre for Flourishing, Universities of Oxford (UK), Aarhus (Denmark) and Pompeu Fabra (Spain); Department of Psychology, University of Amsterdam, Amsterdam, The Netherlands

## Abstract

Ongoing fluctuations in brain state, largely driven by neuromodulators of arousal, shape how sensory inputs are processed and decisions are made. Here, we combined pharmacological manipulations and pupillary proxies of arousal to examine how global cortical state dynamics relate to perceptual sensitivity. We applied the thermodynamics-inspired INSIDEOUT framework to electroencephalography (EEG) recordings from two visual discrimination tasks performed under placebo, atomoxetine (noradrenergic enhancer), and donepezil (cholinergic enhancer) challenges. INSIDEOUT quantifies the strength of functional hierarchies within the brain via the computation of temporal irreversibility in neural activity. Trial-wise global irreversibility increased linearly with pupil-indexed arousal and exhibited a nonlinear inverted-U relationship with perceptual sensitivity, while showing no differences across drug conditions. A nonlinear mediation analysis revealed that temporal irreversibility can account for a significant portion of the association between pupil-linked arousal and perceptual sensitivity. At lower arousal, increases in prestimulus irreversibility improved perceptual accuracy, whereas at higher arousal, pre- and poststimulus irreversibility had different effects. These results bridge intrinsic whole-brain dynamics with perceptual decision-making, demonstrating that neuromodulator-linked arousal tunes cortical states and, in turn, perceptual performance. More broadly, the results shed new light on how arousal may reconfigure neural global dynamics.

## Introduction

Spontaneous fluctuations in brain state continuously shape neural and behavioral responses to incoming stimuli, highlighting the tight coupling between intrinsic dynamics and externally-driven information processing. A central driver of these fluctuations is arousal, associated with the activity of a set of distinct neuromodulators, tuning the “settings” of the brain by altering excitability, synaptic strength, and network coupling [1–6]. Beyond local circuit effects, neuromodulation can reorganize the orchestration of large-scale whole-brain dynamics [3,7]. From a dynamical systems perspective, these observations suggest that arousal does not simply scale neural activity but instead shifts the brain between distinct dynamical regimes [5,8–11].

Pupil size, when measured at constant luminance, has become a popular proxy for such arousal state fluctuations, as it has been shown to correlate with the activity of several neuromodulatory sources shaping cortical computation through the release of noradrenaline, dopamine, serotonin and orexin [12–14]. Notably, human and non-human studies have shown that the arousal-performance relationship is not linear but inverted-U-shaped: optimal performance is observed at moderate levels of pupil-linked arousal, with worse performance at lower and higher ends of the arousal spectrum [15–19]. This raises two crucial questions that will be addressed in the present study. First, how does (pupil-linked) arousal tune global brain dynamics, making one brain state better than another for the processing of sensory information? Second, can we link optimality in behavior and decision-making (“peak performance”) to specific regimes of these global dynamics, shedding new light on the mechanistic underpinnings of the relationship between arousal and behavior?

To address these questions, we bridge the gap between whole-brain intrinsic neural dynamics and human perceptual decision-making by leveraging a novel thermodynamics-inspired framework [20,21]. In this study, we use a neural measure derived from this framework that quantifies the temporal irreversibility of brain signals [20]. The temporal irreversibility of a time series is an indirect measure of the presence and strength of functional hierarchies within a system [21–23]. In a system in which information flows in a symmetric manner, any exchange between different nodes can occur with the same probability as its opposite, and the resulting activity is therefore reversible. In contrast, if information flows asymmetrically, activity in some nodes is causal to the activity in others in a hierarchical way that cannot be reciprocated, leading to temporally irreversible system dynamics (Fig 1A). Irreversibility of activity, or the presence of an “arrow of time”, is then a signature of the complex hierarchical organization that is necessary for optimal computation in the brain [24].

**Fig 1.**
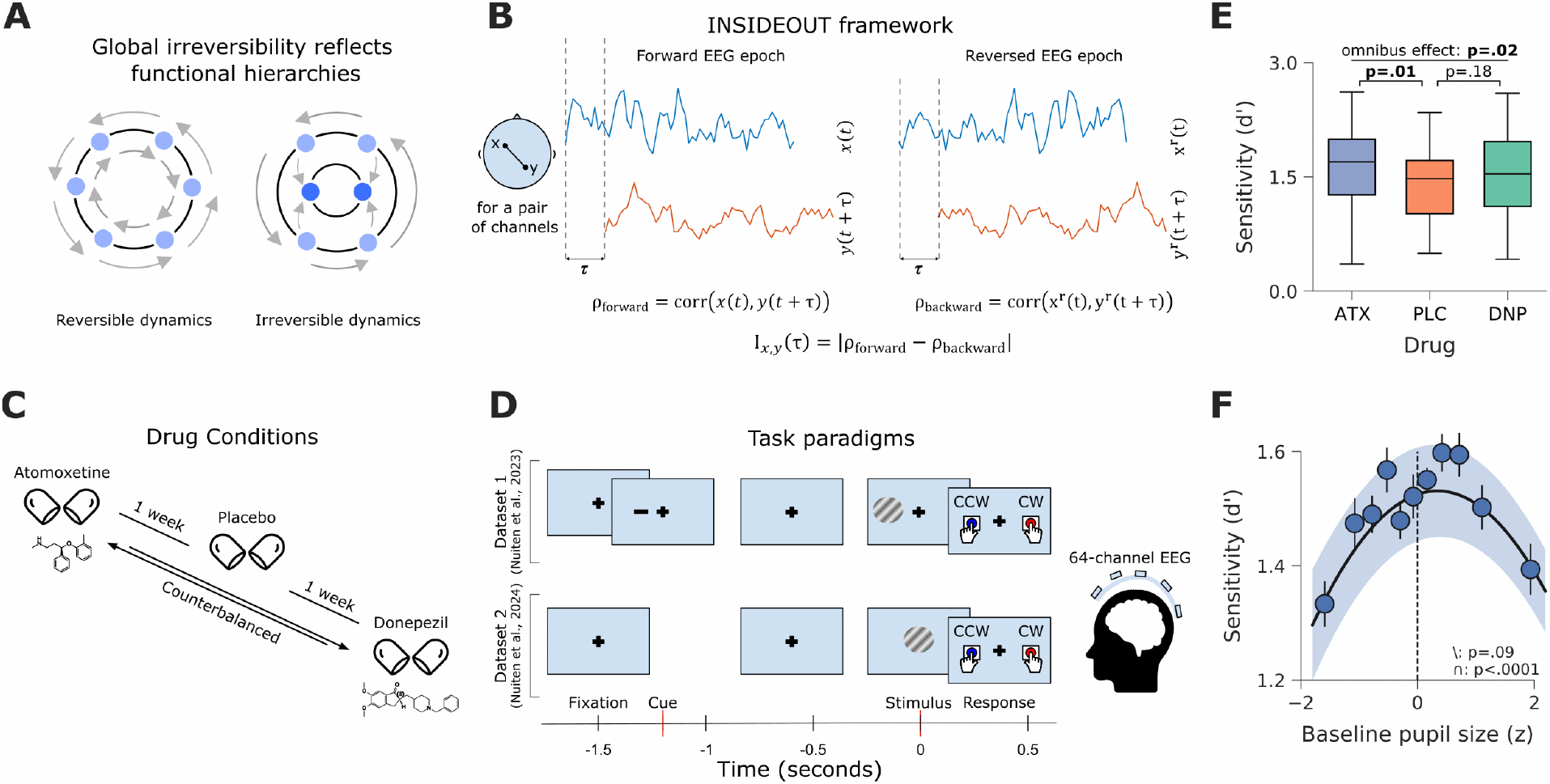
Overview of the methodology and experimental context. (**A**) Global irreversibility value is an indirect measure of the presence of functional hierarchies. In a system that exhibits symmetric information flow (left), there is no arrow of time. On the contrary, when this symmetry of information is broken (right), the system exhibits irreversible dynamics, and the arrow of time has a distinguishable direction. (**B**) The INSIDEOUT framework: pair-wise irreversibility between two time series is computed as the difference between the time-shifted correlation of the forward activity and the time-shifted correlation of its reversed versions. Then, whole-brain global irreversibility is obtained averaging the pairwise irreversibilities of all possible pairs of channels. (**C**) Arousal levels were assessed by measuring pupil size and were manipulated using drugs, administered in counterbalanced sessions approximately one week apart. (**D**) 64-channel EEG data was obtained during two similar visual discrimination tasks: one cued [25] and the other uncued [26]. (**E**) Perceptual sensitivity for different drug conditions: atomoxetine (ATX), placebo (PLC) and donepezil (DNP) across both tasks. Perceptual sensitivity significantly increased in the ATX condition compared to PLC, as shown previously [25,26]. (**F**) Consistent with the Yerkes-Dodson’s law, perceptual sensitivity exhibited an inverted u-shaped relationship with pupil-linked arousal across the two datasets. Shaded area in panel F indicates the standard error of the mean (SEM) over subjects.

There are many ways to assess functional hierarchies in the brain. Some recent efforts have shown that cognitively demanding tasks yield a stronger hierarchical organization compared to rest [27], and that this can be a sensitive marker of neuropsychiatric diseases [28,29]. The INSIDEOUT method is based on quantifying irreversibility of neural activity by comparing the temporal structure of time series in forward time with that of the same signals reversed in time (Fig 1B). Using this approach, neural irreversibility was observed to be higher for conscious than for unconscious states, as well as during demanding task performance compared to rest [20]. This framework outperforms functional connectivity in characterizations of brain states in MEG data [30], and has also elucidated signatures of neurodegenerative disorders when applied to fMRI [31] and EEG data [32]. Extending these advances, we here apply this framework to neural activity recorded during ongoing task performance, quantifying irreversibility in real time rather than only by contrasting “static states”, such as rest versus active or patient versus control groups.

We applied the INSIDEOUT framework to EEG data recorded across three pharmacological conditions: placebo, atomoxetine (driving catecholaminergic tone) and donepezil (driving cholinergic tone; Fig 1C), during two similar visual discrimination tasks (Fig 1D). We explore how changes in spontaneous arousal, indexed by baseline pupil dilation and manipulated with pharmacology, are related to global irreversibility, and how these factors relate to perceptual performance, which is known to be optimal at intermediate arousal states.

## Results

### Perceptual sensitivity increases under elevated catecholaminergic drive and peaks at intermediate pupil-linked arousal

Participants discriminated the clockwise or counter-clockwise orientation of unilaterally (dataset 1 [25]) and centrally presented Gabor patches (dataset 2 [26]), overlaid with dynamic visual noise. Task difficulty was individually titrated to 75% choice accuracy during the intake session and was fixed across the three subsequent pharmacological sessions. Previously, we reported that in both tasks atomoxetine, but not donepezil, increased perceptual sensitivity (*d*′ [33]) when compared to placebo [25,26] and that trial-by-trial variations in baseline pupil-linked arousal exhibited an inverted-U relationship with *d*′ [15,16,26]. Here, we combined the data from both tasks to increase statistical power for our INSIDEOUT analysis.

To summarize the behavior, a 3×2 (drug x task) repeated measures (rm)ANOVAs revealed a main effect of drug condition on *d*′ (*F*_2,52_ = 4.46, *p =* .02) and post hoc comparisons showed this effect was driven by atomoxetine (atomoxetine vs. placebo: t(26) = 2.62, d = 0.52, p = .01; donepezil vs. placebo: t(26) = 1.38, d = 0.24, p = .18; Fig 1E).

We then re-assessed the relationship between baseline pupil-linked arousal and *d*′ by fitting a mixed-effects probit regression model, which formulates signal detection theory (SDT [33]) as a generalized linear model [34,35]. This implementation of SDT facilitates the estimation of trial-wise effects on *d*′ and eliminates the need to subdivide trials in pupil-based bins, as we previously did, thereby improving robustness. Identical to our earlier work, baseline pupil-linked arousal was calculated as the average pupilsize in the 500 ms preceding stimulus onset and trialwise values were z-scored within subject, session, task and run [15,16,26,35]. The model included linear and quadratic pupil-linked arousal terms to capture nonlinear effects and accounted for subject-level variability by including these predictors as random effects.

Fixed-effect interaction terms between the pupil predictors and task were included to capture between-task differences in the pupil-modulation of *d*′. Fixed effects are shown in detail in S1 Table.

This modelling approach revealed a robust inverted-U relationship between baseline pupil-linked arousal and *d*′ (linear pupil: p = .09; quadratic pupil: p < .0001; **Fig 1F**). Furthermore, *d*′ was increased in the cued task (p < .0001) and exhibited a stronger linear relationship with baseline pupil-linked arousal in the uncued task as compared to the cued task (p < .0001). The quadratic relationship between *d*′ and baseline pupil-linked arousal was not different between tasks (p = .34). Thus, *d*′ is higher under pharmacologically increased catecholaminergic drive and peaks at intermediate levels of baseline pupil-linked arousal.

### Global cortical irreversibility is linearly related to baseline pupil-linked arousal

Next, we investigated whether global cortical dynamics were affected by drug conditions and associated with trial-by-trial variations in baseline pupil-linked arousal. We computed global irreversibility for each trial and electrode pair in two time widows, namely the prestimulus epoch (500 ms before stimulus onset) and the poststimulus epoch corresponding to stimulus processing (500 ms after stimulus onset). The global level of irreversibility for each subject was then obtained by averaging across all electrode pairs. To improve robustness, we computed irreversibility for four distinct values of the time shift (τ = [4, 5, 6, 10]), selected based on the mean autocorrelation function across channels and trials. Optimal time-shifts are defined as the smallest lag where the mean autocorrelation function has already decayed sufficiently from its peak at lag 0 (see mean autocorrelation functions in S1 Fig). Centro-parietal and occipital channels contributed most strongly to global irreversibility and this pattern was preserved across time-windows (spatial topographies are shown in Fig 2A and 2E).

**Fig 2.**
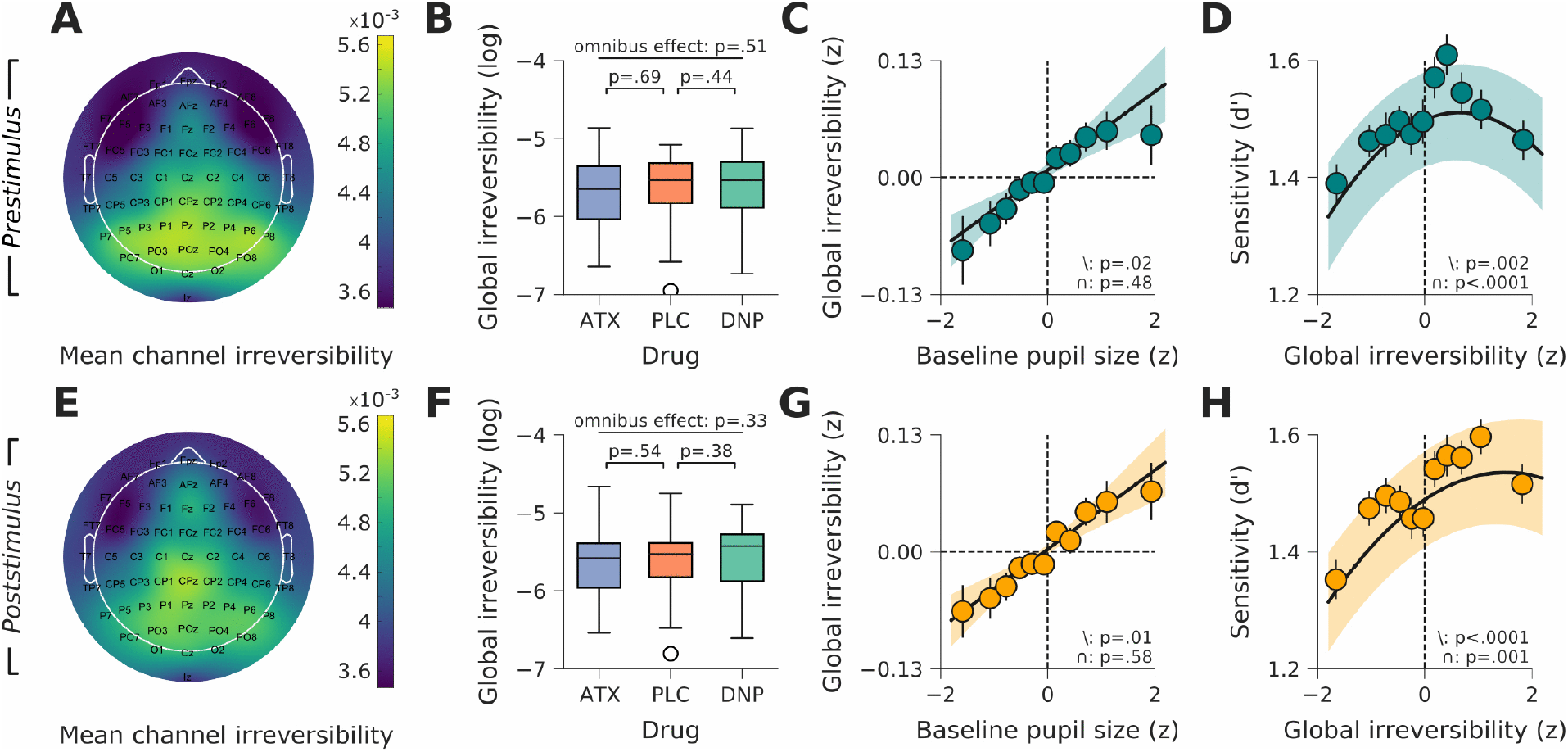
Global irreversibility is associated with pupil-linked arousal and perceptual sensitivity. (**A**) Channel-wise contributions to prestimulus global irreversibility, averaged across subjects, trials, drug condition, and values of τ. (**B**) Prestimulus global irreversibility as a factor of drug condition. (**C**) Prestimulus global irreversibility as a factor of baseline pupil size. (**D**) Perceptual sensitivity (*d*′) as a factor of prestimulus global irreversibility. (**E**-**H**), same as A-D but then for poststimulus global irreversibility. Shaded areas in panels C, D, G, and H indicate the SEM over subjects.

We first assessed whether pharmacological elevation of baseline cholinergic and catechecholaminergic levels, respectively through DNP and ATX, affected global irreversibility. To do so, we log-transformed the irreversibility data to approximate a normal distribution and then averaged irreversibility across trials within subject, drug condition, task, and τ. Irreversibility was then averaged across task and τ before fitting a one-way repeated measures (rm)ANOVA with drug condition as independent variable. We observed no omnibus drug effects for prestimulus (*F*_*2,54*_ = 0.67, p = .51) and poststimulus irreversibility (*F*_*2,54*_ = 1.13, p = .33), and post-hoc pairwise comparisons also did not reveal drug-specific effects (all p > .38; Fig 2B and 2F). Because drug condition did not affect global irreversibility, we did not consider it a factor of interest for all subsequent analyses.

Next, we tested the relationship between baseline pupil-linked arousal and global irreversibility. Log-transformed irreversibility was z-scored within subject, task, and session. We then fitted a linear mixed-effects model in which irreversibility was predicted from baseline pupil-linked arousal. This model included linear and quadratic pupil-linked arousal terms as both fixed and random effects. We further included fixed-effect interaction terms between the pupil-linked arousal predictors and τ and task (see Methods for full model specification). The fixed effects are reported in detail in S2 Table.

Baseline pupil-linked arousal revealed a positive linear relationship with both prestimulus global irreversibility (p = .02; Fig 2C) and poststimulus global irreversibility (p = .01; Fig 2G). Although this linear relationship was stronger in the cued task for both prestimulus global irreversibility (pupil x task interaction: p = .002) and poststimulus global irreversibility (pupil x task interaction: p < .001), marginal slopes revealed that the linear positive effect was observed in both datasets (S2 Fig). The linear relationship between baseline pupil-linked arousal and global irreversibility was also not different across values of τ (all interaction effects p > .18, all marginal slopes p < .02; S2 Fig). We observed no significant quadratic relations between baseline pupil-linked arousal and global irreversibility (prestimulus: p = .48, poststimulus p = .58). Finally, the relation between baseline pupil-linked arousal and irreversibility was spatially-invariant, with occipito-parietal electrodes consistently contributing most to global irreversibility across arousal levels (S3 Fig).

Thus, baseline pupil-linked arousal follows a positive linear relationship with both prestimulus and poststimulus global irreversibility, suggesting that these measures may index a common underlying global neural state rather than independent processes. Because the linear relation between pupil-linked arousal and irreversibility was preserved across the selected range of τ, we decided to average raw irreversibility scores across values of τ for all the subsequent analyses. From here on, we simply refer to this across-τ average as global irreversibility.

### Global irreversibility is nonlinearly related to perceptual sensitivity

Given that baseline pupil-linked arousal follows a positive linear relationship with global irreversibility and an inverted-U relationship with *d*′, global irreversibility may also exhibit an inverted-U relationship with *d*′. To test this, we constructed a similar mixed-effects probit regression model as described before, but now with linear and quadratic irreversibility predictors. Fixed-effect interactions terms between the irreversibility predictors and task were included to account for task-specific effects (Methods, fixed effects are reported in detail in S3 Table).

The model revealed that *d*′ was both linearly and nonlinearly related to prestimulus (linear irreversibility: p = .002; quadratic irreversibility: p < .0001; Fig 2D) and poststimulus global irreversibility (linear irreversibility: p < .0001; quadratic irreversibility: p = .001; Fig 2D). The linear and quadratic relationships between *d*′ and global irreversibility did not differ between tasks for either time-window (S3 Table). Finally, we fitted a model that incorporated both prestimulus and poststimulus irreversibility as predictors. Epoch (prestimulus vs. poststimulus) was included in this model as an additional interaction term with both the linear and quadratic irreversibility predictors to assess whether the relationships between irreversibility and *d*′ were different across time windows. There were no significant differences in either the linear or quadratic predictors between time-windows (S4 Table). Thus, similar to baseline pupil-linked arousal, *d*′ follows an inverted-U relationship to both prestimulus and poststimulus global irreversibility.

Because hierarchical organization in brain networks is closely related to the balance between integration and segregation, which is known to be influenced by neuromodulation [6], we also investigated whether a standard measure of network segregation captures similar relationships with arousal and behavior. We computed modularity, a measure of the extent to which brain activity organizes into segregated subnetworks [36]. Baseline pupil-linked arousal was significantly associated with prestimulus modularity (linear pupil: p = .007, quadratic pupil: p = .008) indicating that arousal modulates the degree of functional segregation in cortical dynamics. However, in contrast to global irreversibility, modularity did not significantly predict *d*′, neither linearly nor quadratically (all p > .30). Critically, when both modularity and global irreversibility were included as predictors in a combined model, irreversibility remained a significant predictor of perceptual sensitivity (linear p = .09, quadratic p = .03), whereas modularity did not (all p > .22). Model comparison further indicated that including modularity did not improve model fit and it was penalized under AIC (AIC of irreversibility model = 31668, AIC of expanded model with modularity = 31681; Δ AIC = −13). In the poststimulus epoch, the same relationship between pupil and modularity was observed, but weaker (linear pupil: p = .06, quadratic pupil: p = .01). In this epoch, modularity showed a significant quadratic relationship with perceptual sensitivity (p = .003). However, again this effect did not explain additional variance beyond irreversibility in combined models (AIC of irreversibility model = 31673, AIC of expanded model with modularity = 31691; Δ AIC = −18), suggesting that the relevance of modularity for predicting *d*′ is not independent of global irreversibility. This indicates that irreversibility captures the behaviorally relevant signature of brain dynamics more effectively than modularity in the context of this study.

As an additional sanity check, we also performed an analogous analysis using the mean absolute activity of each channel. In the prestimulus epoch, it did show an inverted U-shaped relationship with pupil-linked arousal (linear p = .36, quadratic p = .02) but did not predict *d*′ at all (all p > .49). In the poststimulus epoch, the relationship with pupil was best described by a linear relationship instead (linear p = .03, quadratic p = .30), but again it did not predict *d*′ (all p > .32). This analysis allows us to rule out that the observed behavioral effects could be explained by global changes in activity magnitude.

### Global irreversibility mediates relationship between pupil size and choice accuracy

So far, we have established that pupil-linked arousal and global irreversibility are linearly related and that both measures exhibit nonlinear relationships with *d*′ (perceptual sensitivity). These findings may mean that pupil-linked arousal and global irreversibility reflect the same underlying state or may jointly shape behavioral performance (both explaining unique variance). To formally test this, we used statistical mediation modeling. Mediation models quantify how much of the relationship between a predictor (*X*; pupil-linked arousal) and an outcome (*Y*; choice accuracy) is statistically associated with a third variable, referred to as the mediator (*M*; global irreversibility). This approach decomposes the total effect of arousal on choice accuracy into an indirect effect (linked to global irreversibility) and a direct effect (independent of global irreversibility), providing insight into whether global irreversibility contributes to the arousal-behavior relationship. Note that this approach remains fully correlational.

To accommodate for nonlinearities, we constructed a nonlinear mediation model in which the relationships among all three variables were defined as quadratic functions (Methods). In standard linear mediation, the mediator *M* is modelled as a linear function of *X* and outcome *Y* as a linear function of *M*, so that *a*_1_ captures the change in *M* per unit change in *X*, and *b*_1_ captures the change in *Y* per unit change in *M* [37]. Under this formulation, a unit increase in *X* produces an indirect effect of *a*_1_ · *b*_1_ on *Y* through its induced change in *M*. In other words, the mediated effect is defined as the product of the partial derivatives of the functions that govern *X* → *M* and *M* → *Y* [38]. This formulation generalizes to mediation models in which the relationships between predictor, mediator, and outcome contain nonlinearities [38].

In our nonlinear mediation model, the mediated effect was therefore not a scalar value but instead a function of *X*. We defined the instantaneous indirect effect (IIE) as the product of the relevant partial derivatives evaluated at each value of *X* (Methods; Fig 3A). The model was fitted on data for each participant and task, separately for prestimulus and poststimulus irreversibility as mediating variables. Below, we report across-task average model outcomes, task-specific outcomes are reported in S4 Fig.

**Fig 3.**
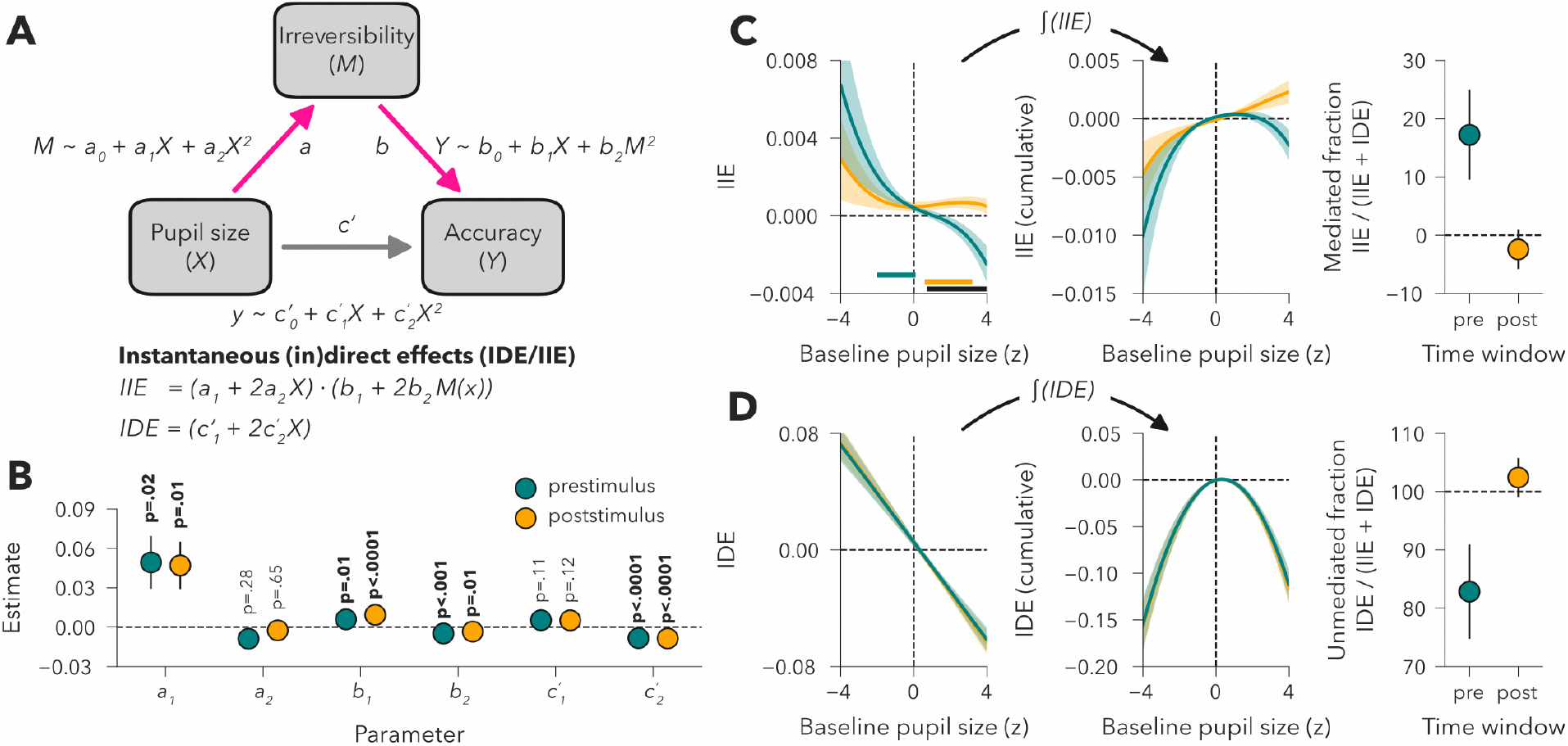
The relationship between pupil-linked arousal and choice accuracy is mediated by changes in global irreversibility. (**A**) Schematic of the nonlinear mediation model. All paths were defined as second-order polynomials. The mediated effect was quantified as the instantaneous indirect effect (IIE), which is defined as the product of the partial derivatives of the *a*-path and *b*-path (expressed in terms of the predictor, *X*). The instantaneous direct effect (IDE) reflects the unmediated effect. Models were fit per task, separately for prestimulus and poststimulus global irreversibility as the mediator. Parameter estimates were averaged across tasks for further analysis. (**B**) Parameter estimates for the linear (demarked with subscript 1) and quadratic (demarked with subscript 2) path components, shown for prestimulus (in teal) and poststimulus (in orange) global irreversibility. (**C**) Left panel: instantaneous indirect effect of baseline pupil size on choice accuracy via changes in irreversibility, shown separately for prestimulus and poststimulus irreversibility. Horizontal bars indicate significant cluster-corrected values of pupil-linked arousal in which mediation was significantly different from 0 (in teal for prestimulus and in orange for poststimulus) or different between the prestimulus and poststimulus model (in black). Middle panel: Cumulative indirect effects, obtained by integrating over the full range of pupil sizes. Right panel: Mediated fraction of the total effect. Shown for prestimulus and poststimulus irreversibility, averaged across their respective clusters shown in the left panel. (**D**) Same as panel C, but for the instantaneous direct effect. Shaded areas indicate SEM.

Consistent with the results shown in Fig 2, pupil-linked arousal displayed a positive linear relationship with both prestimulus and poststimulus global irreversibility (*a*_1_) and a negative quadratic relationship with choice accuracy (*c*′_2_; Fig 3B). Moreover, both prestimulus and poststimulus global irreversibility displayed positive linear and negative quadratic relationships with choice accuracy (*b*_1_ and *b*_2_; Fig 3B). Next, we computed the partial derivatives of *M* as a function of *X* and of *Y* as a function of *M*, expressed in terms of *X*. Multiplying these partial derivatives at each value of *X* yielded the IIE (Fig 3C – left panel). The instantaneous indirect effects for both prestimulus and poststimulus irreversibility revealed significant mediation (assessed with cluster-corrected permutation tests, see Methods), but in distinct pupil-linked arousal regimes (Fig 3C – left panel).

The IIE for prestimulus irreversibility was positive for pupil-linked arousal between −2.0 and +0.1 SD, indicating that increases in pupil-linked arousal within this range improved choice accuracy via changes in prestimulus global irreversibility. Integrating the IIE over the full pupil range indeed revealed an inverted-U shape, in line with the overall relation between pupil size and accuracy (Fig 3C – middle panel). We next calculated the mediated fraction of the total effect to assess the relative contribution of global irreversibility to the overall relationship between pupil-linked arousal and choice accuracy [39]. The total effect is defined as the sum of the IIE and the instantaneous direct effect (IDE), i.e., the derivative of the *c*-path (Fig 3A). The mediated fraction of the total effect is defined as 100 x IIE/(IIE+IDE). Note that this fraction may be negative or exceed 100% when the direct and indirect effects have opposite signs. The mediated effect via changes in prestimulus global irreversibility accounted for 17.22% (± 7.71% standard error of the mean; SEM) of the total effect of pupil-linked arousal on choice accuracy, indicating that the beneficial effect of pupil-linked arousal on behavioral performance at low-to-moderate levels of arousal is partially mediated by prestimulus global cortical dynamics.

Poststimulus irreversibility showed positive IIE values at higher levels of pupil-linked arousal, between + 0.6 and + 3.2 SD (Fig 3C – left panel). Within this elevated pupil-linked arousal range, increases in pupil-linked arousal resulted in improved choice accuracy via changes in poststimulus irreversibility. Integrating the poststimulus IIE revealed an approximate positive linear relation (Fig 3C – middle panel). The mediation effect via poststimulus global irreversibility accounted for −2.43% (± 3.36% SEM) of the total effect of pupil-linked arousal on choice accuracy (Fig 3C – right panel). This negative fraction indicates that, at higher levels of pupil-linked arousal, poststimulus cortical dynamics dampen the detrimental effect of baseline pupil-linked arousal on choice accuracy. This dampening effect is further exemplified by the opposite directions of the IIE and the IDE (shown in Fig 3D) for poststimulus irreversibility at elevated pupil-linked arousal levels.

Importantly, the instantaneous indirect effects of prestimulus and poststimulus irreversibility diverged over a pupil-linked arousal range of +0.7 to +4.0 SD (Fig 3E). In this range, prestimulus effects were more negative, although not significantly below zero, and the poststimulus effect was more positive and mostly significantly greater than zero. This pattern indicates a shift in the locus of mediation, in which higher arousal levels reduce the contribution of prestimulus cortical dynamics to behavior and poststimulus dynamics become the primary mechanism through which arousal shapes choice accuracy.

In summary, both prestimulus and poststimulus global irreversibility mediated the effect of pupil-linked arousal on choice accuracy, indicating that alterations in global cortical dynamics constitute a pathway through which arousal shapes behavior.

## Discussion

Here we asked whether temporal irreversibility of global neural activity tracks fluctuations in arousal and predicts perceptual performance. Pupil-linked arousal fluctuations are associated with the activity of a set of relatively slow-acting neuromodulators (e.g., noradrenaline, dopamine, serotonin) that shape the effects of faster neurotransmitters such as glutamate and γ-amino butyric acid (GABA) [40] (for perception-related pharmacological interventions targeting these latter neurotransmitter systems see for example [41–43]). We found a robust positive linear relationship between baseline pupil size and EEG-based measures of temporal irreversibility. This suggests that spontaneous variations in arousal state are related to the degree to which the brain orchestrates its functional hierarchies, supporting previous work that showed a prominent role of the ascending arousal system in modulating fluctuations in whole-brain cortical dynamics [44,45].

We also investigated the relationship between temporal irreversibility and perceptual sensitivity. We found this relationship to be analogous to the arousal-performance inverted-U relationship, as previously shown [15–17,46]. We add a complementary, whole-brain perspective to this: temporal irreversibility provides a macroscopic index of how arousal-dependent dynamics manifest in large-scale neural activity. Furthermore, by showing that irreversibility scales with baseline pupil size and relates to perceptual sensitivity, we extend the interpretability of this measure beyond distinctly different conditions (e.g., wake vs. anesthesia, rest vs. task) to moment-to-moment fluctuations in brain state during cognition, consistent with the view that functional hierarchies, a signature of non-equilibrium complex brain dynamics [47], are essential for efficient brain function [21]. On top of that, irreversibility captured behavior more sensitively than both a standard measure of network segregation (modularity) and simple mean absolute activity.

Furthermore, building on the findings of how pupil-linked arousal, perceptual sensitivity and neural irreversibility all relate to each other, we tested whether irreversibility statistically mediates the influence of arousal on choice accuracy. A nonlinear mediation model revealed that both prestimulus and poststimulus irreversibility significantly transmitted the effects of arousal onto behavior, but in distinct arousal regimes. At lower baseline arousal levels, increases in arousal improved accuracy via changes in prestimulus irreversibility, whereas at higher baseline arousal levels, poststimulus irreversibility became the dominant route through which arousal enhanced performance. Our results, then, indicate that neural irreversibility reflects a key component of the mechanisms linking arousal to optimal perception, reflecting both spontaneous fluctuations in baseline brain states and stimulus-driven responses.

From a complex systems perspective, the finding that perceptual sensitivity peaks at intermediate levels of neural irreversibility suggests that perception relies on a balance between stability and flexibility in large-scale brain dynamics. A system with flat hierarchies, where activity is highly reversible, may be dominated by noise-like fluctuations around stable attractors: this would limit the effective propagation of information [48]. Conversely, an overly hierarchical system may indicate a brain state with very strong intrinsically driven activity: there already exists a hierarchy of information flow that overrides stimulus-evoked responses. In other words, high irreversibility (where performance drops again after peaking) could reflect a brain state in which a subject got distracted by their own thoughts. Therefore, intermediate levels of irreversibility could be where the dynamical regime operates with an adequate balance between structure and flexibility for integration and processing of stimuli. Together, these findings suggest that arousal (neuromodulation) shapes perception not simply by altering neural gain in local circuits, but by steering whole-brain cortical dynamics toward regimes that are optimally poised for information processing.

## Methods

Participants, task design, and EEG and pupillometry data acquisition and preprocessing steps for this dataset are described in extensive detail in [15,25,26].

### Participants

30 healthy male participants, aged 18-30, were recruited from an online research posting at University of Amsterdam. The study was approved by local ethics and medical committees. Participants underwent extensive screening and assessments to ensure mental and physical health. Written informed consent was obtained from all participants. Two participants withdrew after the first experimental session for personal reasons. Following the initial publication of dataset 2 [15], we excluded one participant who scored near-ceiling on the uncued discrimination task across all sessions and had a *d*′ score > 2.5 SD from the sample mean. To ensure that both datasets were symmetrical in the participants, we also excluded this participant from dataset 1, leading to a final dataset of N = 27.

### Task design

The data we used in this study is the compilation of two different datasets with very similar tasks, which the same subjects completed one after the other in the same sessions. Participants completed three experimental sessions, one for each drug (40 mg atomoxetine, 5 mg donepezil, and placebo), counterbalanced between participants and scheduled at least 1 week apart. Exact administration schedules, physiological effects, and side effects are described in detail in [25]. Both tasks involved discriminating the orientation (clockwise or counterclockwise) of a Gabor patch hidden in dynamic noise (see Fig 1D). In dataset 1 [25], participants’ covert attention was manipulated by an attentional cue (left or right) presented for 300 ms. The attentional cue was predictive of the stimulus location 80% of the time, allowing the separation of trials into validly/invalidly cued groups. After the attentional cue, (and a delay of 1000 ms), the stimulus was then presented to either the left or right side of the screen for 200 ms. Subjects then had to respond with the keyboard to the direction of stimulus orientation. The cue location, stimulus location, and stimulus orientation were all balanced across trials. In dataset 2, the task was slightly different: there was no cue, participants reported the orientation of a centrally presented Gabor stimulus [26].

### Data acquisition and preprocessing

EEG recordings were re-referenced to the average of two earlobe electrodes. A high-pass filter with a cut-off frequency of 0.01 Hz was applied. Automatic detection of faulty EEG channels (via PyPrep toolbox using the RANSAC algorithm [49]), followed by interpolation of faulty EEG channels. Epochs were created from −2 s to 2 s, locked to stimulus onset at 0 ms. Independent Components Analysis (ICA) was used to remove eye-blink artifacts. Another high-pass filter (> 1 Hz) was used on a copy of the epoched data. ICA was fitted to the filtered data with 25 components and components that correlate to EOG electrodes were removed from the epoched data. Artifacts were further removed via the autoreject toolbox [50]. A current source density was computed to minimize volume conductance.

### Measuring irreversibility of brain signals

In this framework [20], irreversibility is quantitatively measured as the amount of asymmetry between the pairwise causal relationships of the forward and reversed time series, computed through the difference between the time-shifted correlation of the forward signals and the time-shifted correlation of their artificially reversed version. The reasoning behind this is that when activity of one time-series is causal to the activity of other time-series course, there should be a certain amount of correlation between these two signals that decays at a certain rate depending on a time shift τ. If causality is directional temporally (presence of an arrow of time), the time-shifted correlation will be lower and/or decay faster for the reversed time-series. Let us consider a multi-dimensional signal *x*(*t*), such as 64-channel EEG data in the case of this study. Let’s consider the time series of channels *i* and *j, x*_*i*_(*t*) and *x*_*j*_(*t*), and their respective reversed versions 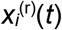 and 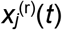.The causal dependency between them is measured through the time-shifted correlation at a given shift τ = *T* (see Fig 1A), expressed as its mutual information to ensure well-behaved, positive values. In the forward case, this is:

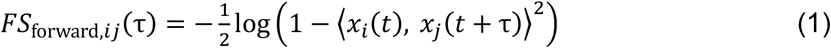

And the backwards case:

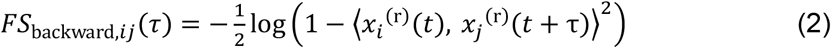

That is, this yields two matrices of time-shifted correlations, one for the forward time series and another one for the reversed time series. Then, the pair-wise irreversibility is given by the quadratic distance between the forward and backwards time-shifted correlation:

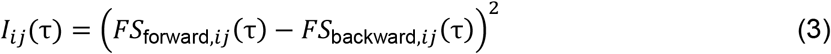

The global level of irreversibility for each subject is obtained by averaging across all elements of ***I***(τ). Raw irreversibility values are nonnegative by definition. However, they follow an extremely right-skewed distribution, so in this study we applied a logarithmic transformation to them to bring their distribution closer to normality.

### Nonlinear mediation model

Before fitting the nonlinear mediation model, we averaged irreversibility across the four τ’s and log-transformed these average irreversibility scores. Next, we z-scored pupil-linked arousal (within subject, task, session and block) and log-transformed irreversibility (within subject, task, and session). The nonlinear mediation model consisted of three paths: the *a*-path describing the relationship between pupil-linked arousal and global irreversibility, the *b*-path describing the relationship between global irreversibility and choice accuracy, and the *c′*-path describing the unmediated (direct) relationship between pupil-linked arousal and choice accuracy. The mediated (indirect) effect is conceived as the change in choice accuracy as a function of pupil-linked arousal via the *a*-path and *b*-path.

All paths were defined as second-order polynomials:

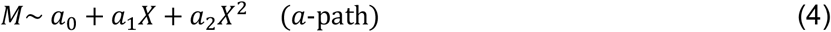

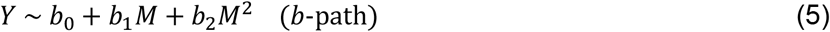

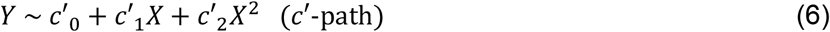

The model was fit per participant and task using Lavaan in R [51]. After fitting, model parameters were averaged per subject across tasks and subject-level parameter estimates were tested against zero with two-tailed one-sample t-tests.

The mediated effect was quantified as the instantaneous indirect effect (IIE), defined as the product of the partial derivatives of the *a*-path and *b*-path [38]. The partial derivatives were thus defined as:

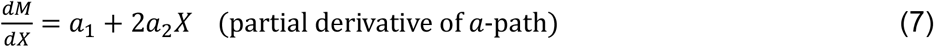

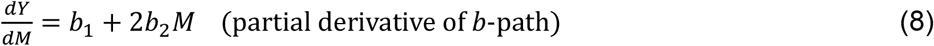

Filling in equation (8) for *M* gives:

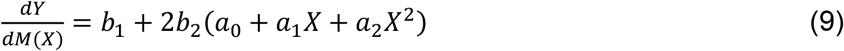

Finally, the IIE was then defined as follows:

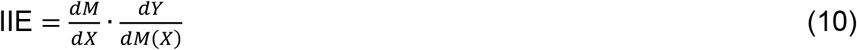

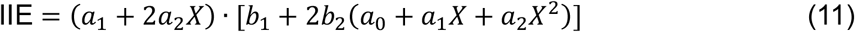

The IIE was calculated over a realistic range of pupil-linked arousal (−4 to +4 SD) in steps of 0.1 SD.

To quantify the strength of the mediation, we calculated the fraction of the total effect accounted for by mediation. The total effect was defined as the sum of instantaneous indirect effect and the instantaneous direct effect (IDE). The mediated fraction was calculated per subject for the pupil size range in which the mediation effects were significant for the respective irreversibility time-windows (prestimulus: −2.0 to +0.1 SD; post-stimulus: +0.6 to +3.2 SD).

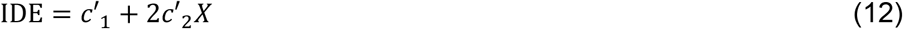

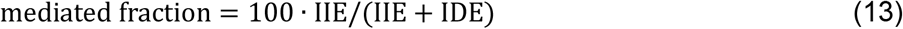

Note that this mediated fraction of the total effect can fall outside of the range 0-100% if the directions of the indirect and direct effects are in opposition.

### Statistical analyses

#### Linear mixed-effects model

To assess the relationship between trial-by-trial variations in baseline pupil size (*X*) and global irreversibility (*I*), we fitted a linear mixed-effects model to pooled data across tasks and values of τ using lmer function of the R package lme4 [52]. The model included fixed effects of baseline pupil size (linear and quadratic terms) and their interactions with task and τ. The model further accounted for inter-subject variability in pupil-modulations of irreversibility by including random intercepts and random slopes for pupil-related effects. Before model fitting, irreversibility scores were z-scored within subject, session, task, and tau, while baseline pupil sizes were z-scored within subject, session, task, and block. Task was effect-coded as −1 (dataset 1) and +1 (dataset 2) and τ was treated as a categorical factor with τ = 10 as the reference level. The fitted model was defined as:

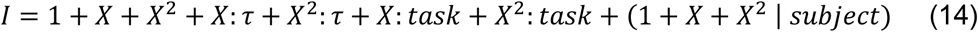

### Mixed-effects probit regression model

To assess the relationships between perceptual sensitivity (*d*′) and trial-by-trial variations in baseline pupil size as well as prestimulus and poststimulus global irreversibility, we used a mixed-effects probit regression model, corresponding to a generalized linear mixed-effects reformulation of SDT [33,34]. Binary responses (*R*; coded as 0 and 1) were modelled using a probit-link function, with stimulus (*S*; effect coded as −0.5 and +0.5) as a predictor such that the stimulus main effect corresponds to SDT *d*′, while the intercept corresponds to response criterion.

Models were fitted using the glmer function from the lme4 in R [52]. Fixed effects included effect-coded stimulus, linear and quadratic terms of z-scored baseline pupil size or global irreversibility (*X, X*^*2*^; either baseline pupil size or global irreversibility), and their interactions with stimulus to capture linear and nonlinear modulation of *d*′ by the predictor. To test whether these relationships differed across tasks, the model further included fixed-effect interactions with task (*T*; effect coded as −1 and +1). Subject-level random intercepts and random slopes were included for stimulus and for stimulus-by-predictor interactions, accounting for inter-individual variability in overall *d*′ and in predictor-related modulation of *d*′.

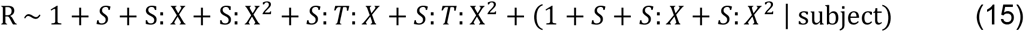

### Mediation model

Nonparametric cluster-based permutation tests against zero (10,000 permutations, cluster threshold: α < .05) were used to evaluate mediation effects over a realistic range of pupil-linked arousal levels (−4 to +4 standard deviations from the mean, step-size of 0.1; SD). Similar cluster-based tests were used to test for difference between mediation for prestimulus and poststimulus irreversibility.

## References

1. Aston-Jones G, Cohen JD. An integrative theory of locus coeruleus-norepinephrine function: adaptive gain and optimal performance. Annu Rev Neurosci. 2005;28: 403–450. doi:10.1146/annurev.neuro.28.061604.135709

2. Ferguson KA, Cardin JA. Mechanisms underlying gain modulation in the cortex. Nat Rev Neurosci. 2020;21: 80–92. doi:10.1038/s41583-019-0253-y

3. Tong C, Li W, Zou Y, Xia Y, Pei M, Zhang K, et al. Norepinephrine-mediated arousal fluctuations drive inverted U-shaped functional connectivity dynamics. Nat Commun. 2025;16: 11318. doi:10.1038/s41467-025-66436-x

4. McCormick DA, Nestvogel DB, He BJ. Neuromodulation of Brain State and Behavior. Annu Rev Neurosci. 2020;43: 391–415. doi:10.1146/annurev-neuro-100219-105424

5. Shine JM. Neuromodulatory control of complex adaptive dynamics in the brain. Interface Focus. 2023;13. doi:10.1098/rsfs.2022.0079

6. Shine JM. Neuromodulatory Influences on Integration and Segregation in the Brain. Trends Cogn Sci. 2019;23: 572–583. doi:10.1016/j.tics.2019.04.002

7. Grimm C, Duss SN, Privitera M, Munn BR, Karalis N, Frässle S, et al. Tonic and burst-like locus coeruleus stimulation distinctly shift network activity across the cortical hierarchy. Nature Neuroscience 2024 27:11. 2024;27: 2167–2177. doi:10.1038/s41593-024-01755-8

8. Guarino D, Carannante I, Destexhe A. Neuromodulators control neuronal dynamics through feature space reshaping. bioRxiv. 2025; 2025.05.19.654470. doi:10.1101/2025.05.19.654470

9. Richman EB, Ticea N, Allen WE, Deisseroth K, Luo L. Neural landscape diffusion resolves conflicts between needs across time. Nature 2023 623:7987. 2023;623: 571–579. doi:10.1038/s41586-023-06715-z

10. Taylor NL, Whyte CJ, Munn BR, Chang C, Lizier JT, Leopold DA, et al. Causal evidence for cholinergic stabilization of attractor landscape dynamics. Cell Rep. 2024;43: 114359. doi:10.1016/j.celrep.2024.114359

11. Johnson PA, Nieuwenhuis S, Mejías J, Urai AE. A dynamical systems model of arousal-driven behavioural state transitions. bioRxiv. 2025; 2025.10.31.685593. doi:10.1101/2025.10.31.685593

12. Joshi S, Gold JI. Pupil Size as a Window on Neural Substrates of Cognition. Trends in Cognitive Sciences. Elsevier Ltd; 2020. pp. 466–480. doi:10.1016/j.tics.2020.03.005

13. Strauch C, Wang CA, Einhäuser W, Van der Stigchel S, Naber M. Pupillometry as an integrated readout of distinct attentional networks. Trends in Neurosciences. Elsevier Ltd; 2022. pp. 635–647. doi:10.1016/j.tins.2022.05.003

14. Cazettes F, Reato D, Morais JP, Renart A, Mainen ZF. Phasic Activation of Dorsal Raphe Serotonergic Neurons Increases Pupil Size. Current Biology. 2020; 1–6. doi:10.1016/j.cub.2020.09.090

15. Beerendonk L, Mejías JF, Nuiten SA, de Gee JW, Fahrenfort JJ, van Gaal S. A disinhibitory circuit mechanism explains a general principle of peak performance during mid-level arousal. Proceedings of the National Academy of Sciences. 2024;121. doi:10.1073/pnas.2312898121

16. Beerendonk L, Mejías JF, Nuiten SA, de Gee JW, Zantvoord JB, Fahrenfort JJ, et al. Adaptive arousal regulation: Pharmacologically shifting the peak of the Yerkes–Dodson curve by catecholaminergic enhancement of arousal. Proceedings of the National Academy of Sciences. 2025;122: e2419733122. doi:10.1073/pnas.2419733122

17. Papadopoulos L, Jo S, Zumwalt K, Wehr M, Jaramillo S, McCormick DA, et al. Modulation of metastable ensemble dynamics explains the inverted-U relationship between tone discriminability and arousal in auditory cortex. Neuron. 2026;114: 740-758.e19. doi:10.1016/j.neuron.2025.11.011

18. McGinley MJ, David SV, McCormick DA. Cortical Membrane Potential Signature of Optimal States for Sensory Signal Detection. Neuron. 2015;87: 179–192. doi:10.1016/j.neuron.2015.05.038

19. Nuiten SA, Lohuis MO, Schaub AC, van Gaal S, Olcese U, Pennartz C, et al. Baseline activity of V1 interneurons connects pupil-linked arousal to engaged behavioral state. bioRxiv. Cold Spring Harbor Laboratory; 2026. p. 2026.02.05.704022. doi:10.64898/2026.02.05.704022

20. Deco G, Sanz Perl Y, Bocaccio H, Tagliazucchi E, Kringelbach ML. The INSIDEOUT framework provides precise signatures of the balance of intrinsic and extrinsic dynamics in brain states. Commun Biol. 2022;5. doi:10.1038/s42003-022-03505-7

21. Kringelbach ML, Sanz Perl Y, Deco G. The Thermodynamics of Mind. Trends Cogn Sci. 2024;28: 568–581. doi:10.1016/j.tics.2024.03.009

22. Deco G, Lynn CW, Sanz Perl Y, Kringelbach ML. Violations of the fluctuation-dissipation theorem reveal distinct nonequilibrium dynamics of brain states. Phys Rev E. 2023;108: 064410. doi:10.1103/PhysRevE.108.064410

23. Nartallo-Kaluarachchi R, Asllani M, Deco G, Kringelbach ML, Goriely A, Lambiotte R. Broken detailed balance and entropy production in directed networks. Phys Rev E. 2024;110: 034313. doi:10.1103/PhysRevE.110.034313

24. Deco G, Vidaurre D, Kringelbach ML. Revisiting the global workspace orchestrating the hierarchical organization of the human brain. Nat Hum Behav. 2021;5: 497–511. doi:10.1038/s41562-020-01003-6

25. Nuiten SA, de Gee JW, Fahrenfort JJ, van Gaal S. Catecholaminergic neuromodulation and selective attention jointly shape perceptual decision making. eLife. 2023. doi:10.7554/eLife.87022.1

26. Nuiten SA, de Gee JW, Zantvoord JB, Fahrenfort JJ, van Gaal S. Pharmacological Elevation of Catecholamine Levels Improves Perceptual Decisions, But Not Metacognitive Insight. eNeuro. 2024;11: ENEURO.0019-24.2024. doi:10.1523/ENEURO.0019-24.2024

27. Kringelbach ML, Sanz Perl Y, Tagliazucchi E, Deco G. Toward naturalistic neuroscience: Mechanisms underlying the flattening of brain hierarchy in movie-watching compared to rest and task. Sci Adv. 2023;9. doi:10.1126/sciadv.ade6049

28. Deco G, Sanz Perl Y, Johnson S, Bourke N, Carhart-Harris RL, Kringelbach ML. Different hierarchical reconfigurations in the brain by psilocybin and escitalopram for depression. Nature Mental Health. 2024;2: 1096–1110. doi:10.1038/s44220-024-00298-y

29. Acero-Pousa I, Escrichs A, Clara Dagnino P, Sanz Perl Y, Kringelbach ML, Uhlhaas PJ, et al. Reconfiguration of functional brain hierarchy in schizophrenia. Transl Psychiatry. 2025;15: 356. doi:10.1038/s41398-025-03584-0

30. Tewarie PKB, Hindriks R, Lai YM, Sotiropoulos SN, Kringelbach M, Deco G. Non-reversibility outperforms functional connectivity in characterisation of brain states in MEG data. Neuroimage. 2023;276: 120186. doi:10.1016/j.neuroimage.2023.120186

31. Sadeghi F, del Agua Banyeres E, Pizzuti A, Okar A, Grimm K, Gerloff C, et al. The arrow of time in Parkinson’s disease. Neuroimage Clin. 2025;47: 103834. doi:10.1016/j.nicl.2025.103834

32. Cruzat J, Herzog R, Prado P, Sanz Perl Y, Gonzalez-Gomez R, Moguilner S, et al. Temporal Irreversibility of Large-Scale Brain Dynamics in Alzheimer’s Disease. The Journal of Neuroscience. 2023;43: 1643–1656. doi:10.1523/JNEUROSCI.1312-22.2022

33. Green DM, Swets JA. Signal detection theory and psychophysics. New York: Wiley. Peninsula Publishing; 1966.

34. DeCarlo LT. Signal Detection Theory and Generalized Linear Models. Psychol Methods. 1998;3. doi:10.1037/1082-989X.3.2.186

35. Nuiten SA, de Gee JW, Zantvoord JB, Sterzer P, Fahrenfort JJ, van Gaal S. Phasic and tonic arousal distinctly shape human decision bias. Commun Biol. 2026;9: 553. doi:10.1038/s42003-026-09776-8

36. Sporns O, Betzel RF. Modular Brain Networks. Annu Rev Psychol. 2016;67: 613–640. doi:10.1146/ANNUREV-PSYCH-122414-033634

37. Hayes AF. Beyond Baron and Kenny: Statistical Mediation Analysis in the New Millennium. Commun Monogr. 2009;76: 408–420. doi:10.1080/03637750903310360

38. Hayes AF, Preacher KJ. Quantifying and Testing Indirect Effects in Simple Mediation Models When the Constituent Paths Are Nonlinear. Multivariate Behav Res. 2010;45: 627–660. doi:10.1080/00273171.2010.498290

39. MacKinnon DP, Warsi G, Dwyer JH. A Simulation Study of Mediated Effect Measures. Multivariate Behav Res. 1995;30: 41–62. doi:10.1207/S15327906MBR3001_3

40. Shine JM. Neuromodulatory Influences on Integration and Segregation in the Brain. Trends Cogn Sci. 2019;23: 572–583. doi:10.1016/j.tics.2019.04.002

41. van Loon AM, Scholte HS, van Gaal S, van der Hoort BJJ, Lamme V a. F. Gaba A agonist Reduces Visual Awareness: A Masking–EEG Experiment. J Cogn Neurosci. 2012;24: 965–974. doi:10.1162/jocn_a_00197

42. Noorman S, Stein T, Zantvoord J, Fahrenfort J, van Gaal S. A causal role of the NMDA receptor in recurrent processing during perceptual integration. Elife. 2025;13. doi:10.7554/elife.100530

43. Noorman S, Fahrenfort JJ, Heilbron M, Sergent C, Zantvoord JB, van Gaal S, et al. Effects of Expectation, Attention, and NMDA Receptor Blockade on Feedforward and Feedback Processing. The Journal of Neuroscience. 2026;46: e0674252026. doi:10.1523/JNEUROSCI.0674-25.2026

44. Munn BR, Müller EJ, Wainstein G, Shine JM. The ascending arousal system shapes neural dynamics to mediate awareness of cognitive states. Nat Commun. 2021;12: 6016. doi:10.1038/s41467-021-26268-x

45. Shine JM, van den Brink RL, Hernaus D, Nieuwenhuis S, Poldrack RA. Catecholaminergic manipulation alters dynamic network topology across cognitive states. Network Neuroscience. 2018;2: 381–396. doi:10.1162/NETN_A_00042

46. de Gee JW, Mridha Z, Hudson M, Shi Y, Ramsaywak H, Smith S, et al. Strategic stabilization of arousal boosts sustained attention. Current Biology. 2024;34: 4114-4128.e6. doi:10.1016/j.cub.2024.07.070

47. Nartallo-Kaluarachchi R, Kringelbach M, Deco G, Lambiotte R, Goriely A. Nonequilibrium physics of brain dynamics. Phys Rep. 2026;1152: 1–43. doi:10.1016/j.physrep.2025.10.003

48. Stikvoort W, Pérez-Ordoyo E, Mindlin I, Escrichs A, Sitt JD, Kringelbach ML, et al. Nonequilibrium brain dynamics elicited as the origin of perturbative complexity. PLoS Comput Biol. 2025;21. doi:10.1371/journal.pcbi.1013150

49. Appelhoff S, Hurst AJ, Lawrence A, Li A, Mantilla Ramos YJ, O’Reilly C, et al. PyPREP: A Python implementation of the preprocessing pipeline (PREP) for EEG data. 2023. doi:10.5281/zenodo.6363575

50. Jas M, Engemann DA, Bekhti Y, Raimondo F, Gramfort A. Autoreject: Automated artifact rejection for MEG and EEG data. Neuroimage. 2017;159: 417–429. doi:10.1016/J.NEUROIMAGE.2017.06.030

51. Rosseel Y. lavaan: An R Package for Structural Equation Modeling. J Stat Softw. 2012;48: 1–36. doi:10.18637/jss.v048.i02

52. Bates D, Mächler M, Bolker BM, Walker SC. Fitting Linear Mixed-Effects Models Using lme4. J Stat Softw. 2015;67: 1–48. doi:10.18637/JSS.V067.I01

