## Supporting Information for "Global nonequilibrium cortical dynamics tie mid-level pupil-linked arousal to optimal task performance in humans"

**Supporting Information** file for manuscript entitled: Global nonequilibrium cortical dynamics tie mid-level pupil-linked arousal to optimal task performance in humans

Elvira del Agua, Stijn Adriaan Nuiten, Jasper B. Zantvoord, Gustavo Deco, Simon van Gaal

| | *Estimate* | *Std. Error* | *z-value* | *p-value* |
|---|---|---|---|---|
| *(Intercept)* | 0.04 | 0.02 | 1.74 | .08 |
| *stimulus* | 1.52 | 0.09 | 17.95 | **$5.03e^{-72}$** |
| *stimulus:pupil_z* | 0.03 | 0.02 | 1.65 | .10 |
| *stimulus:pupil_z_sq* | -0.05 | $7.55e^{-3}$ | -6.91 | **$4.85e^{-12}$** |
| *stimulus:task* | -0.06 | 0.01 | -5.16 | **$2.45e^{-7}$** |
| *stimulus:pupil_z:task* | 0.05 | $9.72e^{-3}$ | 5.45 | **$5.03e^{-8}$** |
| *stimulus:pupil_z_sq:task* | $6.03e^{-3}$ | $6.29e^{-3}$ | 0.96 | .34 |

**S1 Table. Outcomes of the mixed-effects probit regression model, assessing pupil-related modulations of *d′*.** Binary responses were predicted from an intercept (corresponding to signal detection theory (SDT) criterion under effect coding) and effect-coded stimulus ({-0.5, +0.5}, corresponding to SDT *d′*). The model further included two-way interactions between z-scored pupil (linear and quadratic) and stimulus (capturing the linear and quadratic relations between pupil and *d′*), as well as three-way interactions with task as a third variable (capturing task differences in pupil effects on *d′*). Estimates, standard errors, and z-values are shown in scientific notation when the values are smaller than 0.01.

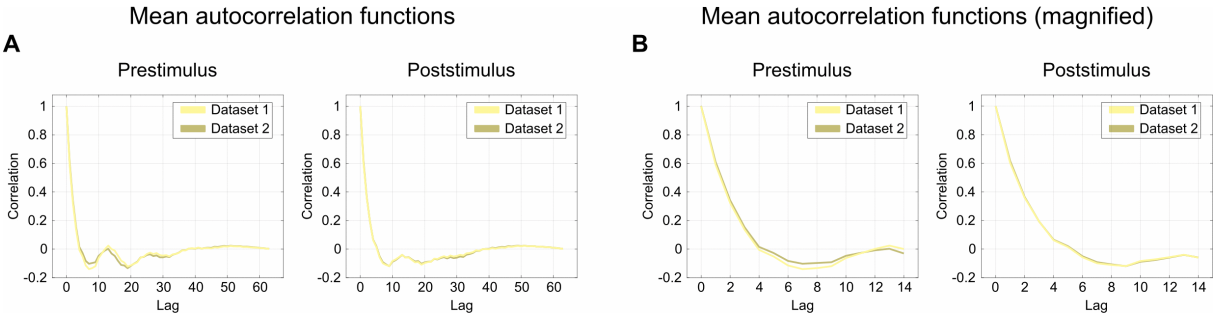


**S1 Fig. Autocorrelation functions over different time-lags.** (**A**) Mean autocorrelation functions for Dataset 1 (in yellow) and Dataset 2 (in khaki) for different time-lags for prestimulus (left panel) and poststimulus (right panel) irreversibility. (**B**) Same as panel A, but magnified.

| | *Estimate* | *Std. Error* | *df* | *t* | *p-value* |
|---|---|---|---|---|---|
| **Prestimulus** | | | | | |
| *(Intercept)* | $8.29e^{-3}$ | $7.31e^{-3}$ | 26.85 | 1.14 | .27 |
| *pupil_z* | $4.36e^{-2}$ | $1.75e^{-2}$ | 28.62 | 2.49 | **.02** |
| *pupil_z_sq* | $5.22e^{-3}$ | $7.33e^{-3}$ | 29.78 | -0.71 | .48 |
| *pupil_z:task* | $-5.45e^{-3}$ | $1.73e^{-3}$ | 354091.15 | -3.15 | **.002** |
| *pupil_z_sq:task* | $7.07e^{-4}$ | $9.52e^{-4}$ | 353680.25 | 0.75 | .45 |
| *pupil_z:tau_4* | $4.25e^{-3}$ | $4.86e^{-3}$ | 354153.98 | 0.88 | .38 |
| *pupil_z:tau_6* | $4.17e^{-4}$ | $4.86e^{-3}$ | 354153.98 | 0.09 | .93 |
| *pupil_z:tau_8* | $6.52e^{-3}$ | $4.86e^{-3}$ | 354153.98 | 1.34 | .18 |
| *pupil_z_sq:tau_4* | $-4.54e^{-3}$ | $2.67e^{-3}$ | 354153.98 | -1.70 | .09 |
| *pupil_z_sq:tau_6* | $-3.10e^{-3}$ | $2.67e^{-3}$ | 354153.98 | -1.16 | .25 |
| *pupil_z_sq:tau_8* | $-4.84e^{-3}$ | $2.67e^{-3}$ | 354153.98 | -1.81 | .07 |
| **Poststimulus** | | | | | |
| *(Intercept)* | $2.90e^{-3}$ | $5.10e^{-3}$ | 27.31 | 0.57 | .57 |
| *pupil_z* | $4.39e^{-2}$ | $1.64e^{-2}$ | 28.83 | 2.67 | **.01** |
| *pupil_z_sq* | $-2.83e^{-3}$ | $5.11e^{-3}$ | 33.51 | -0.55 | .58 |
| *pupil_z:task* | $-6.20e^{-3}$ | $1.73e^{-3}$ | 353475.98 | -3.58 | **$3.45e^{-4}$** |
| *pupil_z_sq:task* | $-2.98e^{-3}$ | $9.53e^{-4}$ | 352384.83 | -3.12 | **.002** |
| *pupil_z:tau_4* | $-3.22e^{-3}$ | $4.87e^{-3}$ | 354154.04 | 0.66 | .51 |
| *pupil_z:tau_6* | $4.58e^{-3}$ | $4.87e^{-3}$ | 354154.04 | 0.94 | .35 |
| *pupil_z:tau_8* | $1.61e^{-3}$ | $4.87e^{-3}$ | 354154.04 | 0.33 | .74 |
| *pupil_z_sq:tau_4* | $-6.09e^{-4}$ | $2.67e^{-3}$ | 354154.04 | 0.23 | .82 |
| *pupil_z_sq:tau_6* | $-5.33e^{-4}$ | $2.67e^{-3}$ | 354154.04 | -0.20 | .84 |
| *pupil_z_sq:tau_8* | $-6.40e^{-5}$ | $2.67e^{-3}$ | 354154.04 | -0.02 | .98 |

**S2 Table. Outcomes of mixed-effects linear model predicting log-transformed irreversibility from trial-by-trial fluctuations in pupil size.** Note that identical standard errors for τ interaction terms reflect the balanced within-subject design and symmetric model specification. Estimates, standard errors, and t-values are shown in scientific notation when the values are smaller than 0.01.

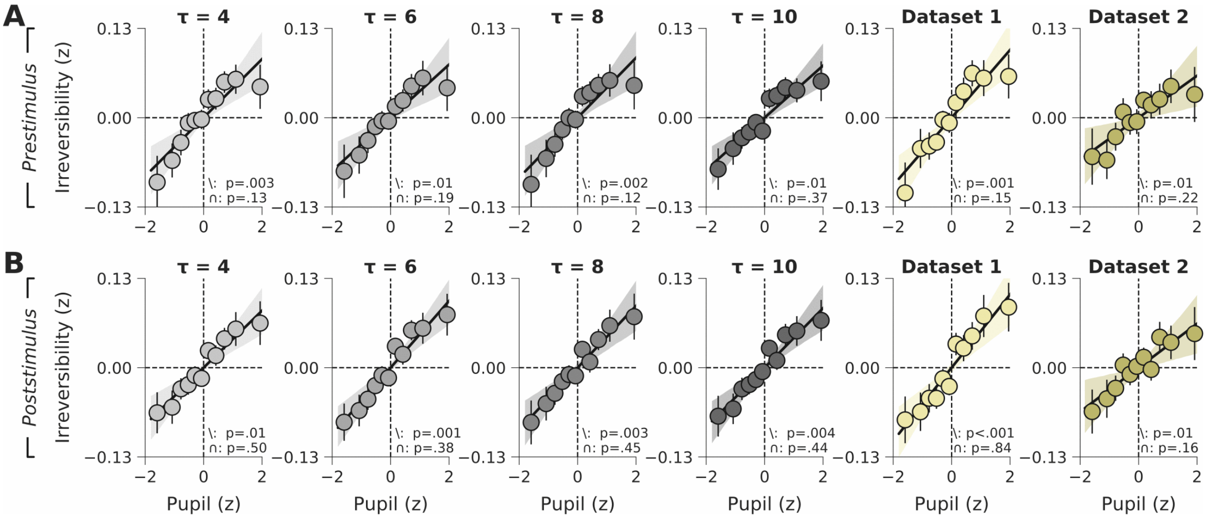


**S2 Fig. Marginalized relationships between pupil-linked arousal and global irreversibility.** (**A**) Marginal trends derived from the mixed-effects model showing the condition-specific effects of pupil-linked arousal on prestimulus global irreversibility, after averaging over other predictors. Across both tasks and all τ values, the linear trend is consistently positive. (**B**) Same as panel A, but for poststimulus global irreversibility. Shaded area and error bars indicate SEM.

| | ***Estimate*** | ***Std. Error*** | ***z*** | ***p*-value** |
|---|---|---|---|---|
| **Prestimulus** | | | | |
| *(Intercept)* | 0.04 | 0.02 | 1.77 | .08 |
| *stimulus* | 1.50 | 0.08 | 18.52 | **$1.42e^{-76}$** |
| *stimulus:irrev_z* | 0.04 | 0.01 | 3.05 | **.002** |
| *stimulus:irrev_z_sq* | -0.03 | $6.96e^{-3}$ | -4.28 | **$1.84e^{-5}$** |
| *stimulus:task* | -0.06 | 0.01 | -5.35 | **$8.96e^{-8}$** |
| *stimulus:irrev_z:task* | 8.90e-5 | $9.47e^{-3}$ | $9.43e^{-3}$ | .99 |
| *stimulus:irrev_z_sq:task* | $5.74e^{-3}$ | $5.61e^{-3}$ | 1.02 | .31 |
| **Poststimulus** | | | | |
| *(Intercept)* | 0.04 | 0.02 | 1.76 | .08 |
| *stimulus* | 1.49 | 0.08 | 18.12 | **$2.21e^{-73}$** |
| *stimulus:irrev_z* | 0.06 | 0.01 | 4.88 | **$1.05e^{-6}$** |
| *stimulus:irrev_z_sq* | -0.02 | $6.28e^{-3}$ | -3.22 | **.001** |
| *stimulus:task* | -0.06 | 0.01 | -5.81 | **$6.12e^{-9}$** |
| *stimulus:irrev_z:task* | -0.01 | $9.463^{-3}$ | -1.22 | .22 |
| *stimulus:irrev_z_sq:task* | 0.01 | $5.48e^{-3}$ | 1.85 | .07 |

**S3 Table. Outcomes of the separate mixed-effects probit regression models, assessing irreversibility-related modulations of *d'*.** Models were separately fit for prestimulus and poststimulus irreversibility. Estimates, standard errors, and z-values are shown in scientific notation when the values are smaller than 0.01.

| | ***Estimate*** | ***Std. Error*** | ***z*** | ***p-value*** |
|---|---|---|---|---|
| *(Intercept)* | 0.04 | 0.02 | 1.77 | .08 |
| *stimulus* | 1.49 | 0.09 | 18.35 | **$3.05e^{-75}$** |
| *stimulus:pupil_z* | 0.05 | 0.02 | 4.61 | **$4.05e^{-6}$** |
| *stimulus:pupil_z_sq* | -0.03 | $6.18e^{-3}$ | -4.01 | **$6.09e^{-5}$** |
| *stimulus:task* | -0.06 | $7.75e^{-3}$ | -7.91 | **$2.61e^{-15}$** |
| *stimulus:window* | $-5.50e^{-3}$ | $7.73e^{-3}$ | -0.71 | .48 |
| *stimulus:pupil_z:task* | $-6.06e^{-3}$ | $6.70e^{-3}$ | -0.90 | .37 |
| *stimulus:pupil_z_sq:task* | $8.04e^{-3}$ | $3.92e^{-3}$ | 2.05 | **.04** |
| *stimulus:pupil_z:window* | 0.01 | $6.68e^{-3}$ | 1.61 | .11 |
| *stimulus:pupil_z_sq:window* | $5.64e^{-3}$ | $3.91e^{-3}$ | 1.44 | .15 |

**S4 Table. Outcomes of the joint mixed-effects probit regression model, assessing irreversibility-related modulations of *d'*.** Both prestimulus and poststimulus irreversibility measures were used as predictors in one model and the epoch time window was included as a fixed-effect interaction term with linear and quadratic irreversibility predictors. Estimates and standard errors are shown in scientific notation when the values are smaller than 0.01.

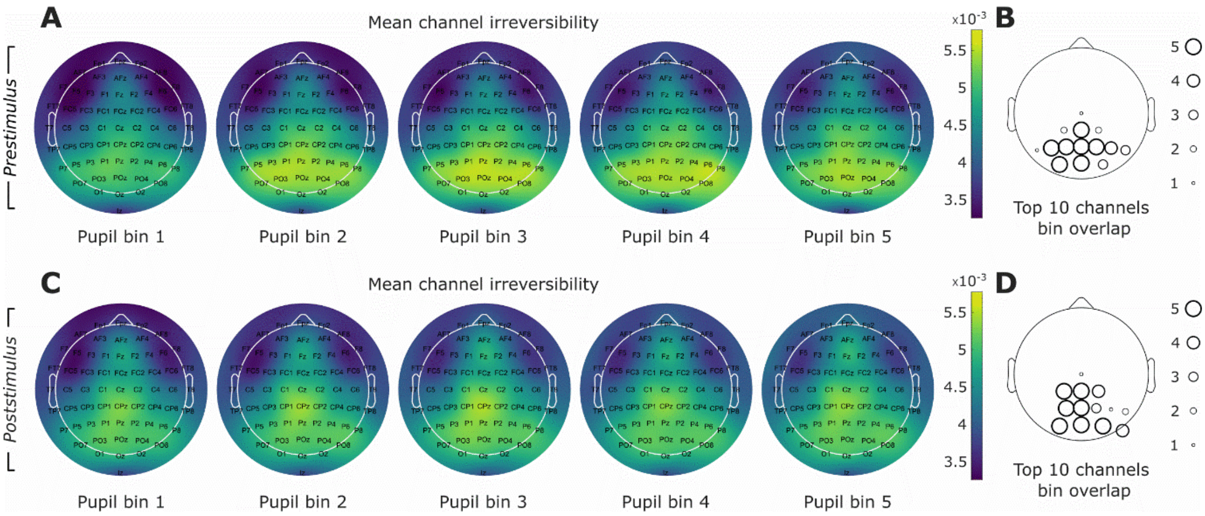


**S3 Fig. Topographic maps of channel contributions to global irreversibility.** (**A**) Topographic maps of channel contributions to global irreversibility during prestimulus epoch and within each pupil bin; averaged across subjects, trials, drug condition and values of values of τ. (**B**) Map displaying the sum of how many times each channel is one of the top 10 contributors to prestimulus global irreversibility within each bin. (**C**-**D**) Same as A-B but for the poststimulus epoch.

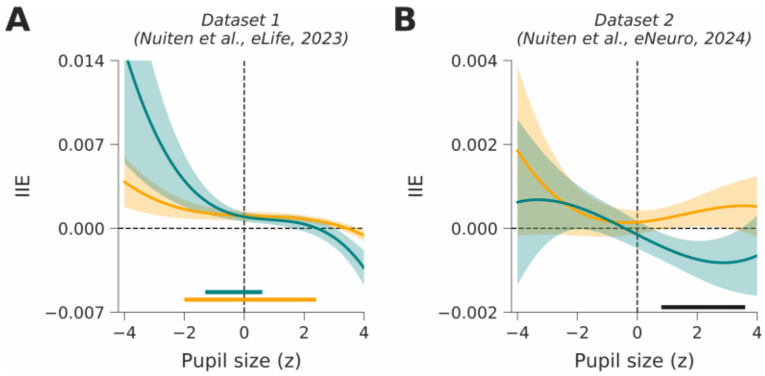


**S4 Fig. Task-specific mediation.** (**A**) The instantaneous indirect effect (IIE) for prestimulus (in teal) and poststimulus (in orange) irreversibility of dataset 1 [25]. (**B**) Same as panel E, but for dataset 2 [26]. Horizontal bars indicate clusters in which IIE was significantly different from zero for prestimulus or poststimulus irreversibility or where prestimulus and poststimulus IIE were different from each other (in black). Shaded areas indicate SEM.
